# Targeted knockdown of *in vitro* candidates does not alter *Wolbachia* density *in vivo*

**DOI:** 10.1101/2025.02.16.638550

**Authors:** Kimberley R. Dainty, Johanna M. Duyvestyn, Heather A. Flores

## Abstract

The bacterial endosymbiont *Wolbachia* has emerged as an effective biocontrol method to reduce arbovirus transmission. Transinfection of *w*Mel *Wolbachia* from *Drosophila melanogaster* to *Aedes aegypti* results in the transfer of important *Wolbachia*-induced phenotypes including the reproductive modification, cytoplasmic incompatibility, and inhibition of viruses including dengue and chikungunya. However, the mechanisms underlying these critical traits as well other *Wolbachia*-host interactions are still not fully understood. Recently an *in vitro* genome wide RNAi screen was performed on *w*Mel-infected Drosophila S2 cells and identified large cohorts of host genes that alter *w*Mel density when targeted. If these findings can be replicated *in vivo*, this would provide a powerful tool for modulating *w*Mel density both systemically and in a tissue-specific manner allowing for interrogation of *w*Mel-host interactions. Here, we used the GAL4/UAS system to express RNAi molecules targeting host gene candidates previously identified to dysregulate *w*Mel density *in vitro*. We found systemic knockdown of two candidate *D. melanogaster* genes does not lead to *w*Mel density dysregulation. To explore the lack of consistency between our study and previous work, we also examined native tissue-specific density of *w*Mel in *D. melanogaster*. We show density is varied between tissues and find that individual tissue densities are not reliable linear predictors of other tissue densities. Our results demonstrate the complexities of implementing *in vitro* findings in systemic applications.

## Introduction

In recent years, the use of *Wolbachia* has emerged as an effective biocontrol tool in multiple mosquito species, including *Aedes aegypti*. Transinfection of the *Wolbachia* strain, *w*Mel, from *Drosophila melanogaster* into *Aedes aegypti* results in the transfer of multiple *w*Mel-induced phenotypes including the reproductive modification of its host (cytoplasmic incompatibility - CI) and the inhibition of medically important arboviruses such as dengue, Zika, and chikungunya (Walker et al. 2011; Hurk et al. 2012; Aliota, Walker, et al. 2016; Aliota, Peinado, et al. 2016; Tan et al. 2017; Carrington et al. 2018; Rocha et al. 2019; Pinto et al. 2021; Utarini et al. 2021). While the *Wolbachia* genes underlying CI have been identified, it is still unclear which host genes and pathways are necessary for CI. Additionally, the virus-inhibiting phenotype induced by *w*Mel and other *Wolbachia* strains is well documented in various *Drosophila* and mosquito species; however, the exact mechanism underpinning it is poorly understood. No study has found a single mechanism of large effect underlying virus blocking (Lindsey et al. 2018). Furthermore, *w*Mel infection in *Ae. aegypti* is also linked to multiple fitness costs, including mild impacts on fecundity, hatch rate, and egg longevity (Fraser et al. 2017; Ant et al. 2018; Allman et al. 2020).

Elucidating the host genes involved in these *w*Mel-host interactions may provide insight into the stability of these critical *w*Mel-induced traits and identify ways to reduce the impact of *w*Mel on host fitness increasing the longevity of this biocontrol method. However, progress in this area has been hindered by the inability to genetically modify *Wolbachia* as well as the inability to easily modulate *Wolbachia* in its hosts.

*In vitro* assays can be a powerful alternative when *in vivo* assays are not feasible. In particular, *in vitro* genome-wide RNAi screens can quickly screen large numbers of genes and identify potential candidate genes involved in a particular pathway or phenotype or interest. In Drosophila, genome-wide screens have been used to successfully identify candidate genes involved in a variety of pathways, including host pathogen interactions (Cherry 2008), virus growth and replication (Cherry 2008), cell growth and viability (Boutros et al. 2004), and RNAi response (Dorner et al. 2006). However, there are also several limitations to *in vitro* genome-wide RNAi screens. The *in vitro* system may not fully recapitulate the complexities of the system (e.g. tissue- or temporal-specific expression) and, of course, there is always the likelihood of off-target effects. Nonetheless, these screens can be a powerful first step to identifying genes and pathways of interest, which can be further interrogated.

Previous work by Grobler et al. (2018) demonstrated that *w*Mel density levels can be dysregulated by way of host gene knockdown *in vitro*. A genome-wide RNAi screen in a *D. melanogaster* cell line revealed large cohorts of host genes that alter *w*Mel density when targeted including an enrichment of ribosomal genes which increased *w*Mel density when downregulated. Two genes associated with ribosomal biogenesis were validated *in vivo* by crossing *w*Mel-infected *D. melanogaster* to mutant allele stocks. As homozygous mutants were lethal, flies that were heterozygous for either ribosomal mutant were used, and heterozygotes showed a 1.2-3-fold increase in *w*Mel densities in germarium and egg chambers when measured by RNA FISH (Grobler et al. 2018). If these findings can be further replicated *in vivo* with the identification of genes which cause both large up and downregulation of *w*Mel density, this would provide a novel way to modulate *w*Mel density within *D. melanogaster*. Combined with additional *Drosophila* tools, this could allow for modulation of *w*Mel both systemically and in a tissue-specific manner, providing a powerful tool to study *Wolbachia*-host interactions.

Here, we used the GAL4/UAS system in *Drosophila melanogaster* to target host genes previously identified to dysregulate *Wolbachia* density when knocked-down *in vitro*. We successfully knocked down two target host genes with RNAi but find no significant changes in *Wolbachia* density. To further explore this, we analysed native tissue-specific density of *w*Mel in *Drosophila melanogaster*. We showed *w*Mel density is not uniform across tissues, and that individual *w*Mel tissue densities are not necessarily predictors of other tissue densities. Our study highlights the complexities of translating *in vitro* findings to a systemic *in vivo* approach.

## Experimental Procedures

### Drosophila melanogaster lines and rearing

*D. melanogaster* lines were obtained from the Bloomington Drosophila Stock Centre, Dept Biology, Indiana University, Bloomington, IN, USA (Supplementary Table 1). For all experiments, flies were maintained on a semolina diet containing, per litre: 7.14 g potassium tartrate, 0.45 g calcium chloride, 4.76 g agar, 10.71 g yeast, 47.62 g dextrose, 23.81 g raw sugar, 59.52 g semolina, 3.56 mL Tegosept, and 3.57 mL propionic acid (Henstridge et al. 2018), at 25°C and ambient humidity.

### RNAi candidate selection

Previously published data by Grobler et al. (2018) was used to identify RNAi candidates to dysregulate *w*Mel *Wolbachia* density. Candidates that had no effect on host cell proliferation (as reported by Grobler et al. (2018)) were ranked by largest degree of *Wolbachia* dysregulation, before being chosen depending on availability of *D. melanogaster* stocks at commercial stock centres with UAS-RNAi transgenes to the genes of interest. Genes chosen had significantly increased *Wolbachia* density with Robust Z scores (average *Wolbachia* per host cell normalised to the plate average) between 2.6 and 5 or had significantly decreased *Wolbachia* density with Robust Z scores between −3.8 and −7.7, as reported by Grobler et al. (2018). Details regarding all *D. melanogaster* lines used in this study can be found in Supplementary Table 1.

### RNAi expression in D. melanogaster

The GAL4/UAS system (Brand and Perrimon 1993) was used to induce expression of the RNAi molecules. The *Tubulin*-GAL4 or *daughterless*-GAL4 promoters were used to drive the ubiquitous expression of RNAi molecules (Supplemental Table 1). Both of these driver strains were infected with *w*Mel-*Wolbachia* by crossing male flies from the GAL4 lines to *w*Mel-infected virgin females from a double balancer (CyO-GFP/IF; TM6B/MKRS) line (Supplementary Fig 1). Virgin females from each GAL4 driver were crossed with UAS-RNAi carrying males once, and progeny assessed for control and desired phenotypes. Control individuals carried the UAS-RNAi transgene, but no GAL4 transgene. Desired individuals carried both the UAS-RNAi and GAL4 transgenes. Control and desired offspring were obtained across multiple replicate vials to obtain enough offspring and minimise vial effects.

### Measurement of RNA expression by reverse transcription (RT) qPCR

For each cross that resulted in viable offspring, both RNA and DNA were extracted from 23-24 control and desired offspring for each cross. Female flies were collected 7 days post emergence, after being allowed to freely mate. For each individual specimen, DNA and RNA were isolated simultaneously. Flies were homogenised in extraction buffer (10 mM Tris pH 8.2, 1mM EDTA, 50 mM NaCl, 25 ug/ml proteinase K), then incubated at 56 °C for 10 minutes, then 37 °C for 30 minutes. DNA and RNA was then extracted using the AllPrep DNA/RNA kit (QIAGEN) following manufacturer’s instructions, with the exception that DNA was eluted in a single 100 µL step. RNA was treated with RNase-Free Dnase Set (QIAGEN) on column. RNA was used to generate cDNA using First Strand cDNA Synthesis kit (K1612, Invitrogen by Thermo Fisher Scientific) according to the manufacturer’s instruction. qRT-PCR was performed using a LightCycler 480 II (Roche) using LightCycler 480 SYBR Green I Master (Roche) according to manufacturer’s protocol. RNA expression was calculated using candidate gene-specific primers relative to the reference *D. melanogaster RpS17* gene (Supplementary Table 2) using the ΔΔCT method (Livak and Schmittgen 2001). RNA expression for each individual knockdown cohort was normalised to the matching individual control cohort.

### Wolbachia density detection by qPCR

The density of *w*Mel *Wolbachia* in desired offspring was compared to that of control offspring from each cross (N=23-24), using DNA extracted as described above. Total relative *Wolbachia* density was determined in whole individual female flies 7 days post emergence (allowed to freely mate), using qPCR with primers to amplify a fragment of the *Wolbachia* gene encoding *wsp*, and the reference *D. melanogaster RpS17* gene (Supplementary Table 2). qPCR was performed for each sample using a LightCycler 480 II (Roche) using the QuantiNova Probe PCR Master Mix (QIAGEN) according to the manufacturer’s protocol. *Wolbachia* density was quantified relative to *RpS17* using the ΔCT method.

### Tissue dissection

7-14-day old female flies (allowed to freely mate) were immobilised on CO_2_ before being dissected in 1x phosphate buffered saline (PBS) solution. Dissected tissues were moved to fresh PBS to rinse away contaminating tissues and haemolymph and processed for DNA isolation.

Individual tissues were homogenised and incubated in extraction buffer (10 mM Tris pH 8.2, 1mM EDTA, 50 mM NaCl, 25 ug/ml proteinase K) at 56 °C and 96 °C for 5 minutes each, before total DNA was isolated from individual tissues using the QIAamp 96 DNA QIAcube HT kit (Qiagen) following manufacturer’s instructions, except in the final step where DNA was eluted in 75 µL of AE buffer. The measurement of *Wolbachia* density in each tissue was performed as described above.

### Statistical analysis

Statistical analysis was performed using GraphPad Prism version 9.0. Data normality was assessed by group using a Shapiro-Wilk normality test. Statistical significance was determined using the Mann Whitney test. Additional statistical analyses and information such as U values, sample sizes and medians can be found in Supplemental Table 3.

## Results

### Host gene knockdown causes no wMel density dysregulation

Grobler et al. (2018) identified 1117 genes that, when targeted with RNAi in cell culture, resulted in a significant increase or decrease in *Wolbachia* density. We chose 14 of these candidates to knockdown to study *in vivo* (six predicted to upregulate and eight predicted to downregulate *w*Mel density). These candidates showed the highest magnitude of *w*Mel density dysregulation, had no effect on host cell proliferation in the Grobler et al. (2018) study, and had RNAi lines available from stock centres. These candidates did not include the two ribosomal biogenesis genes, *RpL27a* and *RpS3*, which were validated *in vivo* by Grobler et al. (2018).

UAS-*RpL27a* could not be sourced, and the magnitude of dysregulation caused by *RpS3* was lower than that of the chosen candidates. Additionally, studies that knocked down *RpS3* with more widely expressed GAL4 drivers resulted in lethality of flies (Perkins et al. 2015).

Transgenic lines containing RNAi molecules targeting candidate genes that dysregulated *w*Mel density *in vitro* were assessed in the GAL4/UAS system using the ubiquitous driver, *Tubulin*-GAL4. Of the 14 crosses performed, seven were found to be lethal under ubiquitous *Tubulin* expression (Figure 1). The crosses were repeated with a driver of lower ubiquitous expression, *daughterless*-GAL4 (Scialo et al. 2016), but found to still be lethal. For the remaining seven crosses where progeny containing both the *Tubulin*-GAL4 and RNAi transgenes were obtained, RT-qPCR was performed. Four of the candidate genes showed no significant change in expression level, one (*dac*) unexpectedly showed significant increase in expression, and two (*CG9801* and *su(r)*) showed a significant decrease in expression compared to controls (Fig 1). We then measured the impact of gene expression modulation on *w*Mel density. The transgenic RNAi strain that caused an unexpected increase in target gene expression (RNAi targeting *dac*) showed a significant increase in *w*Mel density (Fig 2). The candidates that caused significant knockdown of the targeted genes, *CG9801* and *su(r)*, did not cause dysregulation of *w*Mel density (Figure 2). Two other genes showed small, but significant, changes in *w*Mel density, despite no significant change in target gene expression. The transgenic RNAi strain targeting *yippee* showed an increase in *w*Mel density compared to controls, whilst the strain targeting *Pcf11* showed a decrease in *w*Mel density compared to controls.

**Figure 1.**
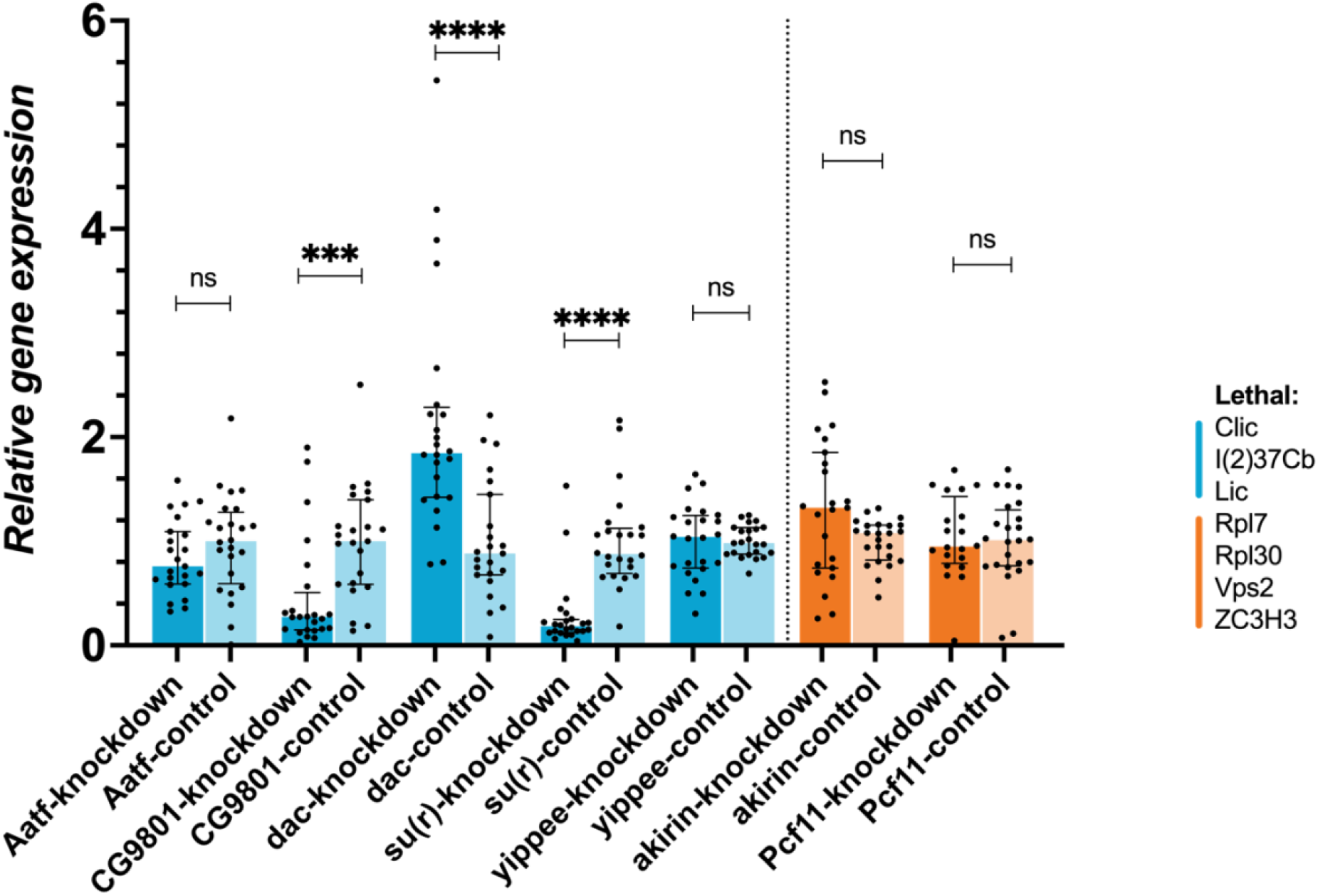
Expression of RNAi-targeted *D. melanogaster* genes. Knockdown of RNAi-targeted genes was assessed using qRT-PCR and significance determined using the ΔΔC_t_ method. Data are the median and IQR of 23-24 flies. Blue bars represent genes identified to decrease *Wolbachia* density when downregulated, and orange bars represent genes identified to increase *Wolbachia* density when downregulated in Grobler et al. (2018). Labels on the X axis indicate genotype of offspring from crosses. Bars labelled ‘knockdown’ represent offspring carrying both the *Tubulin*-GAL4 driver and the UAS-RNAi transgenes. Bars labelled ‘control’ represent offspring carrying only the UAS-RNAi transgene. GAL4-driven RNAi expression found to be lethal for offspring indicated under ‘Lethal’. Statistical significance was determined using the Mann Whitney test comparing each knockdown cohort to its matched control cohort (ns = not significant, \*\*\**P*<0.001, \*\*\*\**P*<0.0001).

**Figure 2.**
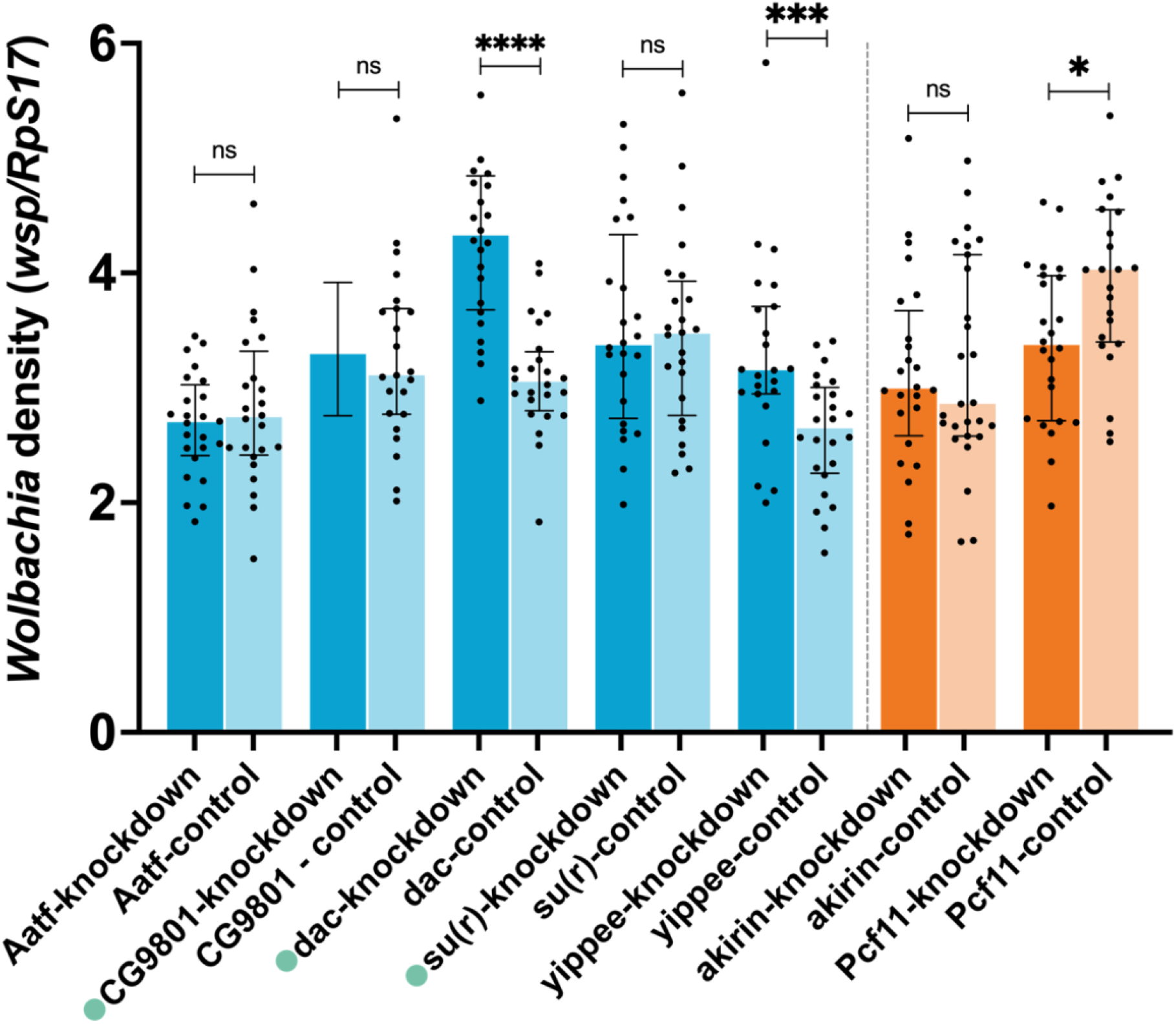
*Wolbachia* density in *D. melanogaster* targeted with RNAi. Density of *w*Mel *Wolbachia* in 7-day old female *D. melanogaster* targeted with RNAi against indicated genes and matched controls were determined by qPCR using primers to amplify a fragment of the *Wolbachia wsp* gene and the reference *D. melanogaster RpS17* gene. The ΔC_t_ method was used to calculate *Wolbachia* density. Data are the median and interquartile range of 23-24 flies. Blue bars represent genes suggested to decrease *Wolbachia* density when downregulated, and orange bars represent genes suggested to increase *Wolbachia* density when downregulated. Labels on the X axis indicate genotype of offspring from crosses. Bars labelled ‘knockdown’ represent offspring carrying both the *Tubulin-*GA4-driver and the UAS-RNAi transgenes. Bars labelled ‘control’ represent offspring carrying only the UAS-RNAi transgene. Bars labelled with a green circle indicate offspring with significant knockdown or upregulation of indicated genes (Figure 1). Statistical significance was determined using the Mann Whitney test comparing each knockdown cohort to its matched control cohort (ns = not significant, \**P*<0.05, \*\*\**P*<0.001, \*\*\*\**P*<0.0001).

### wMel density varies substantially between tissues

One potential explanation for host gene modulation not impacting *w*Mel density systemically, is that *w*Mel density may vary substantially across tissues and successful reduction in some tissues may be overwhelmed by less successful reduction in other tissues. To measure variation in *w*Mel density across tissues, 7-14-day old female flies were collected and the Malpighian tubules, salivary glands, ovaries, fat body, muscles, midgut, and brain were dissected. DNA was isolated from these tissues and *w*Mel density measured by qPCR. *w*Mel density was highly variable across tissue types, with some tissues having nearly one log higher *w*Mel density than others (Fig 3A). The Malpighian tubules and the salivary glands had higher median densities than the other tissues studied. However, the variation in density between individual flies in both tissues was also the largest. Interestingly, flies that had higher density in some tissues did not consistently have high density in other tissues, suggesting that high density is not uniform across tissue types (Fig 3B).

**Figure 3.**
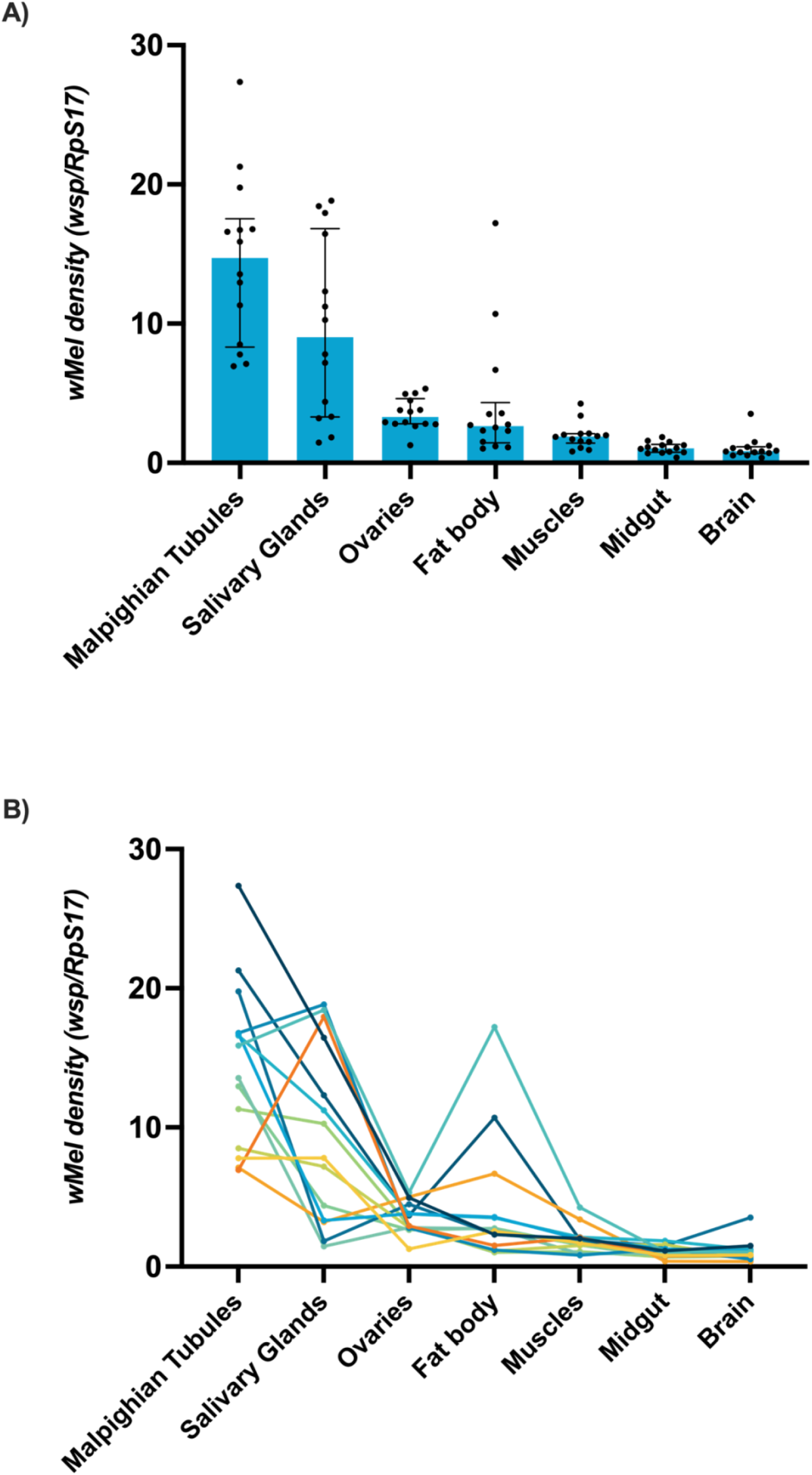
*Wolbachia* density in *D. melanogaster* tissues. Density of *w*Mel *Wolbachia* in 7-14-day old female *D. melanogaster* was determined by qPCR using primers to amplify a fragment of the *Wolbachia wsp* gene and the reference *D. melanogaster RpS17* gene. The ΔC_t_ method was used to calculate *Wolbachia* density. A) Data are the median and interquartile range of 14 flies. B) Each coloured line and associated dot points represent the *w*Mel densities in tissues of an individual *D. melanogaster*.

## Discussion

Many *Wolbachia-*induced phenotypes are influenced by endosymbiont density (Lu et al. 2012; Osborne et al. 2012; Fraser et al. 2017). Thus, having the ability to modulate *Wolbachia* density *in vivo* may provide a powerful tool to study *Wolbachia*-host interactions. We, therefore, attempted to modulate *w*Mel density regulation in *D. melanogaster* genetically. We identified host candidate genes from the Grobler et al. (2018) *in vitro* study to knockdown in an *in vivo* systemic model. RNAi was used in the GAL4/UAS system to target these genes to dysregulate *w*Mel density. Despite our selection of candidates that did not impede cell proliferation *in vitro*, half of the candidates were lethal when ubiquitously knocked-down *in vivo* using two different *Gal4* drivers. Of the remaining candidates, the majority were not significantly knocked-down when targeted with gene-specific RNAi molecules. However, we only assayed one RNAi line per gene. As RNAi lines can show significant variability in their targeting success due to both the RNAi sequence used and the location of the RNAi insertion (Grill et al. 2023), it is possible that if additional RNAi lines were assayed, we would see more successful gene knock down.

Two candidate genes, *CG9801* and *su(r)*, did show significant knockdown. They did not, however, have the same impact systemically in *D. melanogaster* as was observed in the cell line. No perturbation to *Wolbachia* density was seen in the knockdown offspring compared to controls, suggesting ubiquitous targeting of these host genes individually does not systemically affect *Wolbachia* density *in vivo*. As discussed in multiple reviews and studies (Yamamoto-Hino and Goto 2013; Heigwer et al. 2018; Grill et al. 2023), RNAi both *in vitro* and *in vivo* can give rise to false discoveries due to many technical reasons in addition to reasons unique to the biology of *Wolbachia*. First, it is possible that the level of gene knockdown achieved here was lower than that achieved in the Grobler et al., 2018 study, and thus not enough to dysregulate *w*Mel density. As the Grobler et al. 2018 study did not report gene expression levels post targeting, it is not possible to compare between the two studies. Second, it is possible that genes that regulate *w*Mel density in an *in vitro* embryonic cell line (S2 cells) are of lesser importance to *in vivo* due to lower overall expression in adults. This does not seem to be the case for *CG9801* and *su(r)* as its expression in embryonic tissues is similar or higher than its expression in adult tissues (Graveley et al. 2011). Finally, our study also did not assess if ubiquitous expression of the candidate RNAi molecules dysregulated *w*Mel densities variably among tissues. Multiple factors could be at play to cause this. First, expression of the target genes studied here vary across *D. melanogaster* tissues where *w*Mel is present. For example, the expression of *CG9801* and *su(r)* are reported as very high and high respectively in the Malpighian tubules, but low and moderate, respectively, in the salivary glands (Chintapalli et al. 2007). This may have led to *w*Mel dysregulation in some tissues being masked by no or less-dysregulated *w*Mel densities in other tissues. Second, as we have shown here, *w*Mel density varies substantially across different tissues which may interact further with variable host gene expression to mask *w*Mel density modulation. Ideally, host candidate genes with high ubiquitous expression across most tissues would be key targets for downregulation. However, many of these genes likely have important functions (e.g. ribosomal proteins) and their knockdown may be lethal.

Our work here shows that *w*Mel density is highly variable across tissues. *Wolbachia* density variation across tissues has also been observed in other species of *Drosophila* as well as in mosquitoes (Dobson et al. 1999; Osborne et al. 2012; Amuzu et al. 2015; Amuzu and McGraw 2016; Kaur et al. 2020). Moreover, the tissue localisation and density of *Wolbachia* can vary among strains and can change when *Wolbachia* are transinfected into novel hosts (Osborne et al. 2009; Fraser et al. 2017, 2020). The two tissues with the highest *w*Mel density in this study, the Malpighian tubules and salivary glands, have been shown to have high *Wolbachia* density in both *w*Mel-transinfected *Ae. aegypti* as well as other *Drosophila* species (Moreira et al. 2009; Lu et al. 2012; Amuzu et al. 2015; Kaur et al. 2020). One notable between-species difference is the density of *Wolbachia* is consistently higher in mosquito ovaries than in *Drosophila* ovaries when compared to other tissue types (Fraser et al. 2017; Kaur et al. 2020). Interestingly, in the two tissues that presented the highest *w*Mel density here, Malpighian tubules and salivary glands, density was also the most variable. This suggests that *Wolbachia* density may be more tightly regulated in some tissue types than others. Alternatively, *Wolbachia* may be less costly to the host in these tissues and thus be less tightly constrained. For example, it has been suggested that the high density of *Wolbachia* in the Malpighian tubules, excretory organs for insects, is due to the abundance of resources at this site (FARIA and SUCENA 2013) such as electrolytes and organic solutes including amino acids (Beyenbach et al. 2010), which have been shown to be beneficial for *Wolbachia* growth (Caragata et al. 2014). This may indicate that these resources make it less costly to the host for *Wolbachia* to grow to high abundance here or may be more favourable for *Wolbachia* growth.

Interestingly, *w*Mel tissue densities were found to not necessarily be linear predictors of other tissue densities. Individual flies were found to have high *w*Mel densities in particular tissues when compared to other flies, but low *w*Mel densities in other tissues compared to the same flies. Furthermore, individual flies did not always follow the tissue-specific density trends observed across the overall population (Fig. 3). For example, some flies showed higher *w*Mel density in the salivary glands than the Malpighian tubules, or higher density in the fat body than the ovaries, even though the latter tissues had higher median densities across the population. These data showcase the variability of *w*Mel tissue propensities between individuals. Of note, the age of our flies was quite broad (7-14 days), and thus we cannot eliminate the possibility that age may interact with tissue-specific *w*Mel-density. Overall, our study supports previous work suggesting systemic *w*Mel density is not necessarily an appropriate predictor for studies interested in *w*Mel effects or interactions in specific tissues (Osborne et al. 2012; Kaur et al. 2020). These findings should be considered during experimental design.

## Conclusions

In conclusion, we found targeted knockdown of individual host genes found to dysregulate *w*Mel density *in vitro* did not transfer phenotypically *in vivo*. This study depicts the complexities of validating *in vitro* findings *in vivo*. We observed a large amount of lethality caused by gene knockdown despite choosing candidates that did not impact cell proliferation *in vitro* and found that when knocked-down, the majority of target genes did not show a significant decrease in gene expression. For the two genes where knockdown was achieved, no significant impact on *Wolbachia* density was observed, which may be due to the large variation observed in *w*Mel density across tissues. Further work is needed to understand whether individual host gene knockdown can disrupt regulation of *w*Mel density, or whether a more complex multigene approach is needed as the ability to modulate *w*Mel can provide a powerful took to dissect the role of *w*Mel in specific phenotypes. Perhaps focusing specifically on tissue-specific knock-downs in the future will allow for better success in the replication of the *in vitro* observations and/or alleviate issues of lethality associated with candidate gene knock-down.

## Acknowledgements

We would like to acknowledge the contributions of Ritzel Gimeno and Limom Lim for technical assistance. Stocks obtained from the Bloomington Drosophila Stock Center (NIH P40OD018537) were used in this study.

## Notes

### Competing Interest Statement

The authors have declared no competing interest.

