## Supplemental Data for "Targeted knockdown of *in vitro* candidates does not alter *Wolbachia* density *in vivo*"

**Supplementary Data**

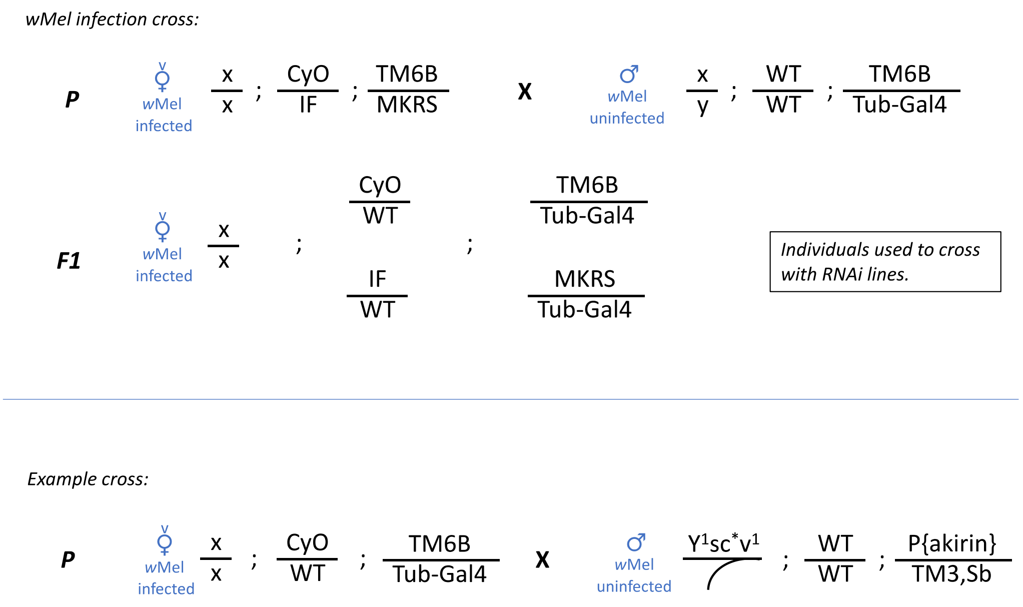


**S1 Figure. Infection of *w*Mel-*Wolbachia* into *Tubulin*-GAL4 and *daughterless*-GAL4 lines and subsequent example use.** *w*Mel-*Wolbachia* was infected into GAL4 driver lines (*Tubulin* and *daughterless*) by crossing males of these lines to virgin females of a double-balancer line (CyO-GFP/IF; TM6B/MKRS) carrying *w*Mel. Virgin females of each GAL4 driver line were then used for subsequent GAL4/UAS crosses.

**S1 Table. *D. melanogaster* lines used in this study.** Columns indicate line name, Bloomington Drosophila Stock Centre (BDSC) Research Resource Identifier (RRID), and genotype of the lines.

| Line name: | BDSC RRID: | Genotype: |
|---|---|---|
| **UAS-Aatf** | BDSC_57211 | y[1] sc[*] v[1]; P{y[+t7.7] v[+t1.8]=TRiP.HMC04594}attP40 |
| **UAS-CG9801** | BDSC_63034 | y[1] v[1]; P{y[+t7.7] v[+t1.8]=TRiP.HMS05308}attP40 |
| **UAS-Clic** | BDSC_66973 | y[1] sc[*] v[1]; P{y[+t7.7] v[+t1.8]=TRiP.HMS05439}attP40/CyO |
| **UAS-dac** | BDSC_35022 | y[1]sc[*] v[1]; P{y[+t7.7] v[+t1.8]=TRiP.HMS01435}attP2 |
| **UAS-I(2)37Cb** | BDSC_56931 | y[1] sc[*] v[1]; P{y[+t7.7] v[+t1.8]=TRiP.HMC04370}attP40 |
| **UAS-lic** | BDSC_60010 | y[1] v[1]; P{y[+t7.7] v[+t1.8]=TRiP.HMS05002}attP40 |
| **UAS-su(r)** | BDSC_53339 | y[1] sc[*] v[1]; P{y[+t7.7] v[+t1.8]=TRiP.HMC03568}attP40 |
| **UAS-Yippee** | BDSC_54020 | y[1] v[1]; P{y[+t7.7] v[+t1.8]=TRiP.HMJ21446}attP40 |
| **UAS-akirin** | BDSC_34036 | y[1] sc[*] v[1]; P{y[+t7.7] v[+t1.8]=TRiP.HMS01010}attP2/TM3,Sb[1] |
| **UAS-Pcf11** | BDSC_32411 | y[1] sc[*] v[1]; P{y[+t7.7] v[+t1.8]=TRiP.HMS00406}attP2 |
| **UAS-Rpl7** | BDSC_34600 | y[1] sc[*] v[1] sev[21]; P{y[+t7.7] v[+t1.8]=TRiP.HMS01074}attP2 |
| **UAS-Rpl30** | BDSC_80379 | y[1] sc[*] v[1]; P{y[+t7.7] v[+t1.8]=TRiP.HMC06615}attP40/CyO |
| **UAS-Vps2** | BDSC_38995 | y[1] sc[*] v[1]; P{y[+t7.7] v[+t1.8]=TRiP.HMS01911}attP40 |
| **UAS-ZC3H3** | BDSC_61816 | y[1] v[1]; P{y[+t7.7] v[+t1.8]=TRiP.HMJ23259}attP40 |
| **da-Gal4** | BDSC_55850 | w[*]; Kr[If-1]/CyO; P{w[+mW.hs]=GAL4-da.G32}UH1 |
| **Tubulin-Gal4** | * | P{w+; Tub-GAL4}/TM3, Sb [1] |

*This line was kindy donated by the Johnson lab, Monash University, Australia.

**S2 Table. Primers and probes used in this study.**

| Target Gene | Forward Primer | Reverse Primer | Probe | Reference |
|---|---|---|---|---|
| ***Aatf*** | GCAATCAGTCCATGTGGTAAAC | CGTCTGTGGCTGTTTCTTTG | - | This study |
| ***CG9801*** | AGCATCCCAGCATCCATTT | CAGCGACGGCATCTTTAGT | - | This study |
| ***dac*** | CAGCGACGGCATCTTTAGT | GGCGCTAACGAGTGCTATTT | - | This study |
| ***su(r)*** | GGACCTGCTGTCGCTAAAT | TGCGTTTCCAGTGCTTCT | - | This study |
| ***yippee*** | GTAGGCACATCTAGCACGTAAG | CTGAGTAGCAGCCCATCTTT | - | This study |
| ***Akirin*** | GATCAAGGAGCGCGAGAAT | ACAAAGGCATCGTACTGCTC | - | This study |
| ***Pcf11*** | GCCAGTCGTTGAGATCGTAATA | CACTTCCGACAGAATCGTAGAG | - | This study |
| ***RpS17*** | AGCGTCGTGACAACTACGTC | TCAACATCTCCTTGGTGTCG | CGTCTCCGCTCTGGAGCA GG | This study |
| ***wsp*** | ATTGAAGATATGCCTATCACTCC | GTTACATCATAACTAACACCAGC | TCCTTTGGAACCCGCTGTG AATGA | (Frentiu et al. 2014) |

**S3 Table. Additional statistical analyses and information of relative gene expression after RNAi targeting, and *Wolbachia* density analysis.**

| | | **Medians** | **U value** | **Samples sizes** | **Significance** |
|---|---|---|---|---|---|
| Relative gene expression | *Aatf*-control vs. *Aatf*-knockdown | 0.7586 \| 1.000 | 226 | 23 \| 24 | 0.2951 |
| | *CG9801*-control vs. *CG9801*-knockdown | 0.2723 \| 1.000 | 117 | 24 \| 23 | 0.0005 |
| | *dac*-control vs. *dac*-knockdown | 1.846 \| 0.8819 | 83 | 24 \| 23 | <0.0001 |
| | *su(r)*-control vs. *su(r)*-knockdown | 0.1833 \| 0.8763 | 48 | 24 \| 24 | <0.0001 |
| | *yippee*-control vs. *yippee*-knockdown | 1.040 \| 0.9831 | 271 | 24 \| 23 | 0.9244 |
| | *akirin*-control vs. *akirin*-knockdown | 1.322 \| 1.076 | 191 | 23 \| 24 | 0.0719 |
| | *Pcf11*-control vs. *Pcf11*-knockdown | 0.9502 \| 1.012 | 236 | 23 \| 24 | 0.9349 |
| *Wolbachia* density | *Aatf*-control vs. *Aatf*-knockdown | 2.701 \| 2.745 | 260 | 23 \| 24 | 0.5739 |
| | *CG9801*-control vs. *CG9801*-knockdown | 3.295 \| 3.110 | 257.5 | 24 \| 23 | 0.7004 |
| | *dac*-control vs. *dac*-knockdown | 4.326 \| 3.054 | 49 | 24 \| 23 | <0.0001 |
| | *su(r)*-control vs. *su(r)*-knockdown | 3.371 \| 3.472 | 283 | 24 \| 24 | 0.9268 |
| | *yippee*-control vs. *yippee*-knockdown | 3.153 \| 2.648 | 124 | 24 \| 23 | 0.0009 |
| | *akirin*-control vs. *akirin*-knockdown | 2.994 \| 2.859 | 320 | 23 \| 24 | 0.9478 |
| | *Pcf11*-control vs. *Pcf11*-knockdown | 3.374 \| 4.030 | 171 | 23 \| 24 | 0.0153 |
